# Δ^9^-Tetrahydrocannabinol (THC) impairs visual working memory performance: A randomized crossover trial

**DOI:** 10.1101/778068

**Authors:** Kirsten C. S. Adam, Manoj K. Doss, Elisa Pabon, Edward K. Vogel, Harriet de Wit

**Affiliations:** Department of Psychology, *University of California San Diego*; Institute for Neural Computation, *University of California San Diego*; Department of Psychiatry and Behavioral Sciences, *Johns Hopkins University*; Department of Psychiatry and Behavioral Neuroscience, *University of Chicago*; Grossman Institute for Neuroscience, Quantitative Biology, and Human Behavior, *University of Chicago*; Department of Psychology, *University of Chicago*; Institute for Mind and Biology, *University of Chicago*

**Author notes:** **Correspondence to:** Kirsten C. S. Adam, University of California San Diego, 9500 Gilman Drive, Mail Code: 0109, La Jolla, CA 92093-0109.

**Keywords:** THC, working memory, mind wandering, metacognition

## Abstract

With the increasing prevalence of legal cannabis use and availability, there is an urgent need to identify cognitive impairments related to its use. It is widely believed that cannabis, or its main psychoactive component Δ^9^-tetrahydrocannabinol (THC), impairs working memory, i.e., the ability to temporarily hold information in mind. However, our review of the literature yielded surprisingly little empirical support for an effect of THC or cannabis on working memory. We thus conducted a study with 3 main goals: (1) quantify the effect of THC on visual working memory in a well-powered sample (2) test the potential role of cognitive effects (mind wandering and metacognition) in disrupting working memory, and (3) demonstrate how insufficient sample size and task duration reduce the likelihood of detecting a drug effect. We conducted two double-blind, randomized crossover experiments in which healthy adults (*N=23, 23*) performed a reliable and validated visual working memory task (the “Discrete Whole-Report task”, 90 trials) after administration of THC (7.5 and/or 15 mg oral) or placebo. We also assessed self-reported ‘mind wandering’ (Exp 1) and metacognitive accuracy about ongoing task performance (Exp 2). THC impaired working memory performance (*d* = .65), increased mind wandering (Exp 1), and decreased metacognitive accuracy about task performance (Exp 2). Thus, our findings indicate that THC does impair visual working memory, and that this impairment may be related to both increased mind-wandering and decreased monitoring of task performance. Finally, we used a down-sampling procedure to illustrate the effects of task length and sample size on power to detect the acute effect of THC on working memory.

## Introduction

Cannabis and its main psychoactive constituent, Δ^9^-tetrahydrocannabinol (THC), are widely believed to impair working memory, the mental workspace used to hold information “in mind” that is needed for everyday behaviors such as driving and problem solving [1,2]. Disruptions to working memory can thus disrupt ongoing behavior and lead to negative outcomes, from the innocuous (e.g., forgetting that your turn-signal is on) to the dire (e.g., hitting the cyclist you forgot was in your blind-spot). Thus, understanding the acute effects of THC on working memory and cognition is of critical importance for public health and safety.

Despite the popular belief that THC impairs working memory, this impairment has been difficult to demonstrate under controlled conditions. Here we first reviewed 40 within-subjects, randomized, placebo-controlled studies that tested the acute effect of THC on working memory performance, and then conducted a controlled laboratory study. Examining published Digit Span studies from 1970 to 2019, we found that more than 70% of 57 study conditions failed to detect an effect of the drug (p > .05). This difficulty in demonstrating the effects of THC on working memory in controlled studies [3–8] is especially notable given the bias toward publishing ‘positive’ results [9–11], which should inflate reports of positive effects in the literature. We hypothesized that the lack of empirical support for impairment of working memory after THC reflects limitations of previous studies related to insufficient sample sizes and trial counts. We present the results of a study that addresses these problems.

Prior studies have predominantly used one of a few canonical working memory tasks in which subjects must remember verbal material such as letters or numbers over a short delay (e.g., “Digit Span”[12]). However, over the past 20 years, tasks that test memory for visual information (e.g., colors, shapes, objects) have emerged as popular and robust measures of working memory functioning that are less susceptible to chunking or other strategic factors that are known to impact verbal working memory measures [13,14]. These visual working memory tasks have been demonstrated to be highly reliable [15,16]*, correlate well with other measures of working memory [19,20], and predict individual differences in general fluid intelligence [19]. Further, these tasks been deployed with a variety of patient populations [21–24], have been extensively examined using EEG and FMRI [25–29], and are simple enough to lend themselves to cross-species translational studies [30]. Here, we used a specific variant of a visual working memory task, the “Discrete Whole-Report” task [17,31], to investigate the acute effects of THC on visual working memory performance. A particular advantage of this task over other highly similar tasks (e.g., partial-report [33,34], and change detection [35,36]) and other more distantly related visual WM tasks (e.g., spatial N-back [32]) is that it provides sensitive trial-by-trial measurements of performance [17,37,38] and is thus well-suited for characterizing deficits in the ability to store information.

We hypothesized that two key factors (sample size and task duration) affect power to detect working memory drug effects in a task-generalizable fashion. To test this hypothesis, we performed simulations in which we tested the effect of sample size and task duration (i.e., number of trials) on power in our visual working memory sample, and we compared the predictions of these simulations with results observed in the literature (specifically, the Digit Span literature). Our simulations focus primarily on understanding the task-generalizable effect of sample size and task duration on power. However, the working memory tasks in the reviewed literature differ in other regards, which we later discuss.

We also examined self-reports of mind-wandering (Exp 1) and metacognition (Exp 2), processes that may be modulated by the acute effect of THC. In Experiment 1, we tested whether a tendency to have one’s mind “off-task” (i.e., “mind wandering” or “zoning out”) [39] co-occurs with decreased working memory performance and the acute administration of THC. Whereas mind wandering and decreased awareness of mind wandering are known to occur during nicotine withdrawal [40] and alcohol intoxication [41], little is known about the effect of THC on mind wandering during ongoing task performance. In Experiment 2, we tested whether poor metacognition (i.e., performance monitoring) likewise co-occurs with behavioral decrements and the acute administration of THC. Prior work has found acute administration of THC decreased performance monitoring in a simple visual attention task [42,43]. Here, we tested if acute administration of THC likewise disrupts performance monitoring during a working memory task using metacognitive accuracy of task performance [44] as an index of performance monitoring.

The current study had three key goals which together test the effect of THC on visual working memory. First, we sought to characterize the effects of THC on visual working memory using a longer than typical task (90 trials versus ∼15 trials) and higher than typical sample size (combined *n* = 46 vs. *n* = ∼15). Second, we tested visual working memory performance in relation to other ongoing cognitive processes, specifically increased mind wandering (Exp 1) and decreased metacognitive accuracy of task performance (Exp 2). Finally, we examined previous studies on the effect of THC on working memory to determine whether previous failures to detect effects could be related to insufficient power. To this end, we combined our literature review with a down-sampling procedure on our own, well-powered sample (achieved power > .99) to determine the consequence of inadequate sample sizes and task lengths on the ability to detect a working memory impairment (*d* = .65).

## Methods

### Participants

Healthy occasional (non-daily) cannabis users, aged 18-35, were recruited for two experiments. Sample sizes were set *a priori* to n = 24 per study. Procedures were approved by the University of Chicago Institutional Review Board, and participants provided written, informed consent. Studies took place within the Human Behavioral Pharmacology lab at the University of Chicago. Participants were screened with a physical examination, an electrocardiogram, and a semi-structured interview by a clinical psychologist. Exclusion criteria included any current Axis I DSM-IV disorder including substance dependence, current use of >5 tobacco cigarettes per day, history of psychosis or mania, less than a high school education, lack of English fluency, a body mass index outside 19-33 kg/m2, high blood pressure (>140/90), abnormal electrocardiogram, daily use of any medication other than birth control, pregnancy, or lactating. Cannabis use was assessed in an in-person interview. Inclusion criteria for Experiment 1 were lifetime use between 4 and 100 times and non-daily use, and for Experiment 2 some lifetime use but not daily use. Equal numbers of men and women participated in both Experiments, and their mean ages were 23.0 years (SD = 3.6) in Experiment 1 and 23.4 years (SD = 4.3) in Experiment 2. Data from one subject in each study were excluded because of extreme values (>3 SD’s below the mean).

### Drug

THC (Marinol®; Solvay Pharmaceuticals) was placed in opaque, size 00 capsules with dextrose filler. Placebo capsules contained only dextrose (0 mg THC). Chosen doses produce reliable subjective and cardiovascular effects without adverse effects [45,46]. Experiment 1: Participants received a placebo capsule and a 15 mg capsule in two randomized, counterbalanced sessions. Experiment 2: Participants received a placebo capsule, a 7.5 mg capsule, and a 15 mg capsule in three randomized, counterbalanced sessions.

### Design

Subjects participated in two (Experiment 1) or three (Experiment 2) sessions, conducted at least 1 week apart, in a comfortable laboratory setting. Both experiments used double-blind, within-subjects, counterbalanced designs. A lab member that knew the design of the study but was not running participants performed randomization (via a computerized list shuffle method) and sorted placebo and THC capsules into bags (i.e., bags labeled “first capsule and “second capsule” in Experiment 1; bags labeled “first capsule”, “second capsule”, and “third capsule” in Experiment 2).

### Procedures

#### Pre-session

During an orientation session subjects received instructions, signed a consent form, and practiced the tasks. They were instructed to consume their normal amount of caffeine and nicotine, but to abstain from alcohol, prescription drugs (except contraceptives), over-the-counter-drugs, cannabis, and other illicit drugs for at least 48 hours before session. Participants were informed that they would be tested for recent drug use at the beginning of each session, and positive tests would result in rescheduling or dismissal. Finally, they were advised to get their normal amounts of sleep and to not eat for 2 hours prior to each experimental session. To minimize expectancy effects, participants were informed that they may receive a stimulant, sedative, cannabinoid, or placebo during the sessions.

#### Experimental sessions

At the beginning of each laboratory visit, subjects provided breath and urine samples for breath alcohol level (Alco-sensor III, Intoximeters, St. Louis, MO), a urine drug test (ToxCup, Branan Medical Co., Irvine, CA), and a pregnancy test (females only; Aimstrip, Craig Medical, Vista, CA). Those testing positive were rescheduled or dropped from the study. Baseline cardiovascular and mood measures were taken, then participants consumed the capsule (placebo or THC, double-blind). During the first 120 min, participants relaxed with magazines and music while the drug was absorbed. Cardiovascular and mood measures were taken at regular intervals (every 30-60 minutes) throughout the session. Cognitive testing was conducted from 120 to 220 minutes post-capsule. These time points fall in the time range where subjective and behavioral effects of the drug have reached their peak and remain elevated [47]. The key cognitive test of interest was the Discrete Whole Report task (see section “Visual Working Memory Task”, [17,26,31]). During the Whole-Report Task, participants also reported ratings of their level of mind wandering (Experiment 1) or metacognition (Experiment 2). In Experiment 1 the working memory task was performed at around 160 min post-capsule (M = 159.4, SD = 14.5, Range = [126,193]) and in Experiment 2 it was performed 220 min post-capsule (exact time not recorded). In both experiments, subjects also completed other tasks that are reported elsewhere [48–50], and were not expected to interfere with the task reported here.

### Physiological and Subjective Measures

In both experiments, heart rate and blood pressure were recorded at regular time points (every ∼30 minutes, SI Methods) with portable monitors (Experiment 1: A&D Medical/Life Source, San Jose, CA; Experiment 2: Omron 10 Plus, Omron Healthcare). Self-report measures of the drug effects were obtained at the same times. These have been reported elsewhere, and included the Addiction Research Center Inventory [ARCI] [51,52], the Visual Analog Scales [VAS] [53], the Drug Effects Questionnaire [DEQ] [54], and an End of Session Questionnaire (ESQ; Experiment 1 only). The results of all subjective measures are reported in the main text and/or SI Results, and descriptions of the subjective measures are given in the SI Methods.

### Visual Working Memory Task

The visual working memory (“Discrete Whole Report”) task consisted of 90 trials (3 blocks of 30) [17,31]. On each trial, participants briefly viewed (200 ms) an array of 6 brightly colored squares, and remembered the colors and locations of the squares across a delay with a blank screen (1,000 ms), see Fig 1. Colors for each trial were chosen without replacement from a set of 9 highly-discriminable colors [17]. At test, “response grids” appeared at each location (3×3 grid of all 9 colors). Participants freely recalled the color-location pairing of each item by clicking the color in each response grid that corresponded to the color remembered at that location. They were required to make a response to all 6 squares before moving on to the next trial. In both experiments, participants also provided self-reported measures about their performance throughout the task (“Task-Unrelated Thoughts” or “Item-level Confidence Judgements”).

**Fig 1.**
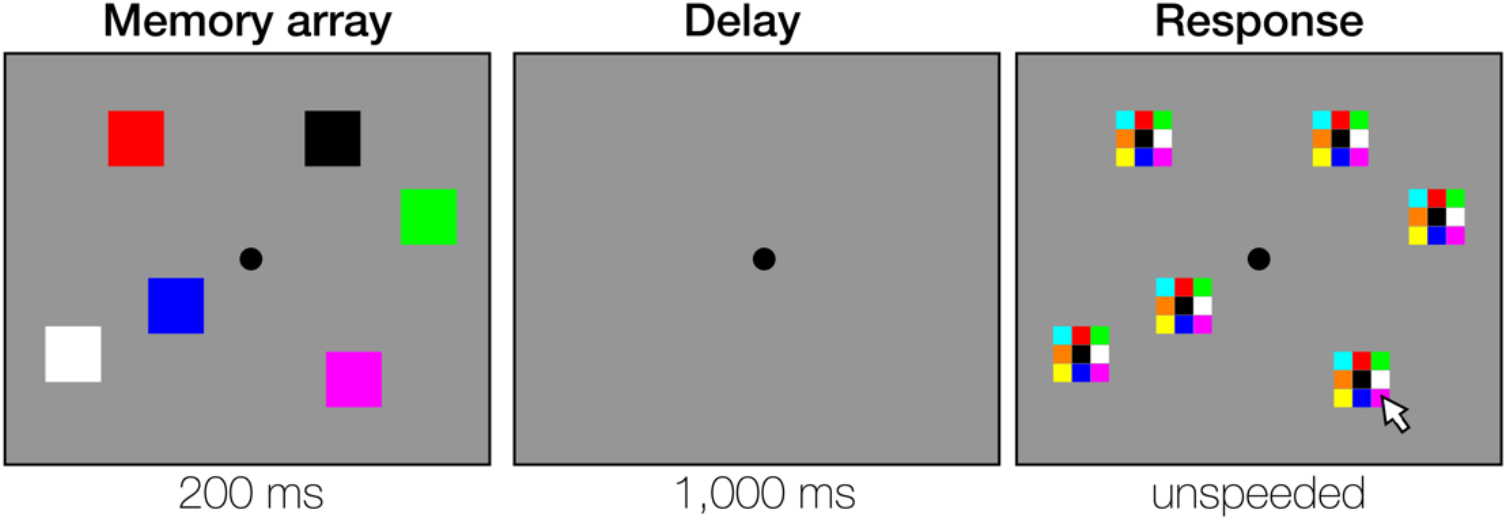
Stimuli used in the whole-report task. On each trial, participants briefly view a memory array containing 6 colored squares (Memory array). Participants remember the colors across a 1,000 ms delay, and report them at test (Response). Participants may report the colors of the items in any order that they choose, and they must make a response for all 6 locations. In the figure, the participant is clicking the magenta subsection of the response grid to indicate that the remembered square in the bottom right corner was magenta.

#### Task-Unrelated Thoughts

In Experiment 1, participants were asked, on 20% of trials, about the contents of their thoughts ‘at the moment’, choosing between three categories: “on task”, “mind wandering”, or “zoning out”. Participants were given instructions and examples of each category during the orientation pre-session. In the instructions, the categories were defined as follows: (1) “on task” indicates that the subject was focused on the task at hand, (2) “mind wandering” indicates that the subject was internally focused on something other than the task, and (3) “zoning out” indicates that the subject was withdrawn and not allocated to anything in particular. If subjects endorsed “mind wandering”, they were asked to classify whether their mind had wandered toward the future, the past, “other”, or “I don’t know”. Although the terms “mind wandering” and “zoning out” are sometimes used interchangeably (e.g. [40]), here we distinguished between periods of internally directed attention (“mind wandering”) and periods where there is a complete absence of attention to anything in particular (“zoning out”, also referred to as “mind blanking”, [55]). The zoning out rating thus differentiates an increase in internally directed attention from a total disengagement of attention.

#### Item-level Confidence Judgements

In Experiment 2, in addition to reporting the color, participants made a binary confidence judgement for each response. While making their responses about the color of each object, they indicated confidence by clicking on the chosen color with either the left or right mouse button. If they felt they had “some information” in mind about the color of the item they were reporting, they should click the color with the left mouse button. If they felt they had “no information” in mind about the color of the item they were reporting, they should click the color with the right mouse button. The number of confident items was calculated for each trial by summing the number of left click responses (ranging from 0 to 6 confident responses per trial). To calculate metacognitive accuracy, we correlated the number of confident responses per trial with the number of accurate responses per trial (e.g., number of items where the correct color was chosen on each trial, ranging from 0 to 6 correct items per trial).

### Statistical Analysis

Data analysis was performed using JASP (Version 0.11.1)[56] and custom scripts in MATLAB 2018A (The MathWorks, Natick, Massachusetts, USA). Datasets for all experiments are available on the Open Science Framework at **https://osf.io/5heur/**. To assess subjective and physiological measures at the time of the working memory test, we calculated a change score from baseline (timepoint closest to the working memory test minus the timepoint immediately before consuming the capsule). In Experiment 1, One participants’ heart rate and blood pressure could not be collected due to device malfunction, leaving 22 participants, and one participants’ subjective measures could not be collected due to a computer malfunction, leaving 22 participants. Significance of placebo versus THC was tested by paired *t*-test (2-tailed) in Experiment 1 (placebo versus 15 mg THC) and tested by 1-way Repeated Measures ANOVA in Experiment 2 (within-subjects factor Drug containing 3 levels, Placebo, 7.5 mg THC, and 15 mg THC).

### Literature Review and Power Analysis

We reviewed the literature to find within-subjects, randomized, placebo-controlled studies testing the acute effect of THC on working memory performance (SI Methods). By far the most common test of working memory was the Digit Span (Forward/Backward) Task. We found 15 papers meeting our inclusion criteria that reported the results of a standard Digit Span task [45,57–70]. Together, these papers reported a total of 57 different conditions that were tested (e.g. Forward vs. Backward span, differing doses of THC). We did not include conditions from papers that reported only combined Digit Span (i.e., forward and reverse not separately reported) [71], or conditions measuring Digit Recall instead of Span [72–76]. These and other tasks show consistent patterns (SI Results), but we chose to focus on only Digit Span conditions for our core arguments because this task (1) is the single most-used task and (2) is administered in a highly consistent manner. See SI Results for the largely non-significant p-values across conditions for other working measures in the literature.

## Results

### Demographic information, subjective measures, and physiological measures

Demographic characteristics of participants are shown in Table 1. Mean values for each of the physiological and subjective measures are shown in Tables S4 and S5 in the supplemental materials.

**Table 1.** Demographic data for Experiments 1 and 2.

|  | Experiment 1 | Experiment 2 |
| --- | --- | --- |
| *Sex* | 12 female; 11 male | 12 female; 11 male |
| *Age (years)* | 22.70 (.72) | 23.52 (.90) |
| *Education (years)* | 15.13 (.35) | 15.57 (.33) |
| *BMI* | 24.77 (.76) | 23.31 (.43) |
| *Caffeine (cups/day)* | 1.43 (.28); n = 22 | 1.69 (0.22); n = 21 |
| *Nicotine (cigarettes/day in the five users)* | .19 (.07); n = 5 | 2.80 (0.85); n = 5 |
| *Alcohol (drinks/week)* | 6.57 (1.38); n = 21 | 8.61 (1.65); n = 21 |
| *Cannabis (uses/month)* | 2.11 (.35); n = 9 | 7.67 (1.86); n = 13 |
| *Lifetime uses of Cannabis* | 27.13 (5.27); n = 23 | 173.25 (250.04); n = 20 |
| *Last use of cannabis before Placebo session (days)* | 118.61 (41.48); n = 23 | 7.82 (1.82); n = 17 |
| *Last use of cannabis before THC session (days)* | 116.20 (41.48); n = 23 | 6.76 (.98); n = 17<br>10.67 (1.69); n = 18 |

This table includes only participants whose data was not excluded for outlier performance (n = 23 per experiment). Age, education, and past month recent substance use are listed as mean (SEM). For recent substance use, the mean and SEM were calculated using only subjects who reported any recent use of the drug (n for each drug type is shown). Two subjects in Exp 2 had missing data for the lifetime use of Cannabis question and one participant was excluded for reporting an outlier value (10,000 uses, >40 SD's outside the other participants' mean number of uses). Remaining participants reported no recent use of the drugs.

THC produced its expected effects on physiological and subjective measures. THC (15 mg) increased heart rate, including at the time of the working memory test in both Exp 1, *t*(21) = 3.55, *p* = .002, *d* = .76, and Exp 2, *F*(2,44) = 12.16, *p* < .001. Only 1 dose (15 mg) was used in Exp 1. In Exp 2 (15 mg, 7.5 mg), there was a linear effect of dose (*p* < .001), but only the high dose was significantly different from placebo (high vs. placebo *p* < .001, low vs. placebo *p* = .25). The drug did not affect systolic or diastolic blood pressure (p > .3). See SI Results for tables of all values. THC also increased scores on the “marijuana scale” of the Addiction Research Center Inventory [ARCI] (*p* < .001), and the “Feel” (*p* < .001), “Like” (*p* ≤ .005), “Dislike” (*p* ≤ .03), and “High” (*p* < .001) questions of the Drug Effects Questionnaire [DEQ] (Bonferroni-corrected for the 5 DEQ measures). The drug increased “Want more” ratings of the DEQ (*p* = .002; *p* = .197), and Visual Analog Scale [VAS] measures “Sociable” and “Friendly” (*p* < .05, Bonferroni-corrected for 13 VAS measures) in Exp 1, but not Exp 2. See SI Results for tables of all values.

### Mean working memory performance

THC impaired working memory performance relative to placebo in both Exp 1 (Fig 2A) and Exp 2 (Fig 2B). In Exp 1, participants correctly reported an average of 3.11 (SD = .49) items in the placebo condition and 2.77 (SD = .50) items in the 15 mg THC condition, *t*(22) = 3.72, *p* = .001, *d* = .78. In Exp 2, participants correctly reported an average of 3.02 (SD = .53) items in the placebo condition, 2.84 (SD = .44) items in the 7.5 mg THC condition, and 2.78 (SD = .54) items in the 15 mg THC condition, *F*(1.52,33.42) = 4.58, *p* = .026, η_p_^2^ = .17^†^. Although polynomial contrasts revealed a linear effect of dose in Exp 2 (*p* = .005), only the high dose was significantly different from placebo (placebo vs. high, *p* = .018; placebo vs. low, *p* = .07).

**Fig 2.**
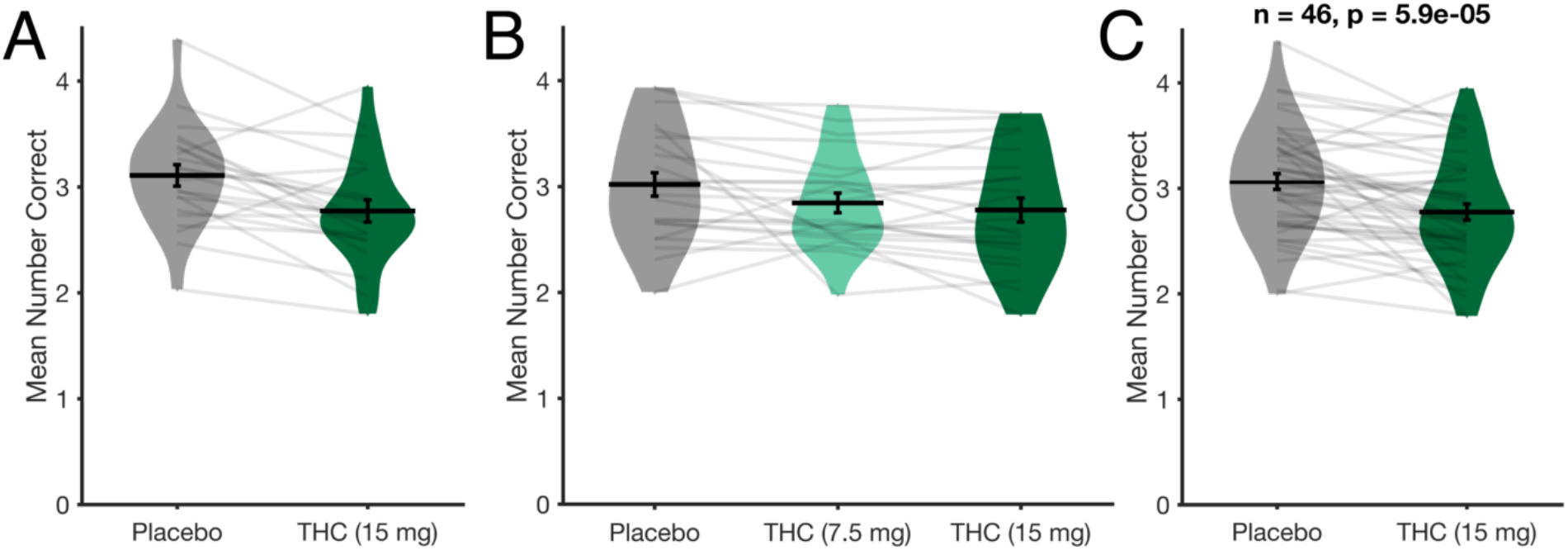
Mean working memory performance. Mean values for the number of items correctly identified in Exp 1 (A; N=23), Exp 2 (B; N=23) and the two experiments combined (C; N=46). Here and elsewhere, violin plots show the distribution of participants, black error bars represent 1 SEM, and transparent gray lines show individual participants. THC significantly reduced the number of remembered items in the 15 mg conditions.

In a separate analysis, we combined the working memory performance data from Experiments 1 and 2. A unique set of participants was recruited for Experiments 1 and 2 so this pooling did not result in multiple data-points from the same participants. Because only Experiment 2 had the 7.5 mg THC condition, this condition was discarded from pooled analyses. We observed the same main effect of THC on working memory performance (Comparing the combined placebo condition to the combined 15 mg THC condition; Fig 2C). A mixed ANOVA with within-subjects factor Drug and between-subjects factor Experiment revealed no main effect of Experiment, *F*(1,44) = .09, *p* = .76, η_p_^2^ = .002, and no interaction between Drug and Experiment, *F*(1,44) = .52, *p* = .48, η_p_^2^ = .01, so this combination of experiments is justified. This combination of experiments yields a total sample size of 46 subjects and a robust effect of Drug on working memory performance, Fig 2C, *t*(45) = 4.43, *p* < .001, *d* = .65. With this larger sample size, we quantified reliability, the effect of experimental block, and the effects of response number on both accuracy and response time. Task reliability (even-odd correlation) was excellent during both the placebo (*r* = .91) and the THC (*r* = .90) conditions, and individual differences in performance were preserved across the THC and Placebo conditions, as shown by a positive correlation (Fig 3A; *r* = .63, *p* < .001, 95% CI [.41, .78]).

**Fig 3.**
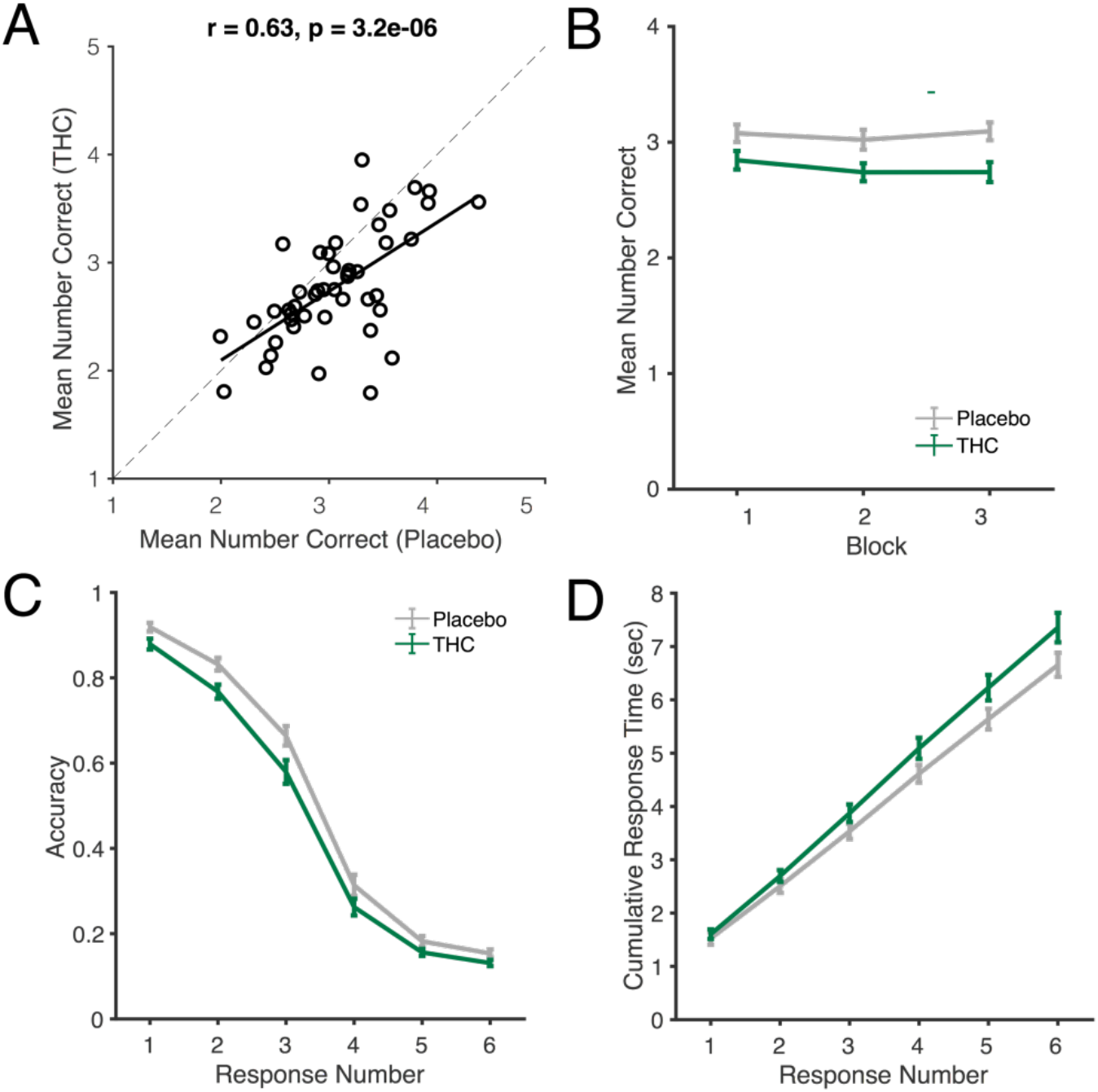
Illustrations of the effects of THC (15 mg) vs placebo on working memory performance (N=46, Exp 1 and Exp 2 combined). (A) Correlation between mean number correct on the working memory task after placebo and THC (15 mg) conditions. This shows that individual differences are reliable across the drug and placebo conditions, but that most individuals are impaired by the drug. (B) Mean number of items correctly identified during the three blocks of the task after placebo and THC (15 mg). Performance was consistently poorer after THC across all three blocks. (C) Working memory performance as a function of response number and drug. Responses were overall less accurate for THC versus placebo, particularly early in the trial. (D) Cumulative response time as a function of response number and drug indicates that impaired performance was not due to a speed-accuracy tradeoff; participants were overall slower for THC versus placebo.

### Changes in working memory performance across experimental blocks and individual responses

To determine whether the effect of THC on working memory performance was related to a decline in effort or task engagement over the course of the experiment, we compared performance across the three blocks of the experiment (30 trials per block). The difference between THC and placebo scores was constant over time (Fig 3B). A repeated measures ANOVA with factors Drug and Block revealed no main effect of Block, *F*(2,90) = 2.69, *p* = .074, η_p_^2^ = .06, and no interaction between Drug and Block, *F*(1.73,77.98) = 1.60, *p* = .210, η_p_^2^ = .03.

To determine whether the effect of THC on working memory was related to careless responding and a speed-accuracy trade-off, we examined response time and accuracy for each response in the trial. If participants were simply more careless at responding in the THC condition, then they may have responded quickly and with poor accuracy. Thus, if the THC-related working memory decrement is driven by a speed-accuracy tradeoff, we should observe faster response times for trials where participants showed lower accuracy. The empirical data did not support a speed-accuracy tradeoff account. Accuracy was overall lower in the Drug condition, particularly for the first three responses. There was a main effect of Drug, *F*(1,45) = 19.64, *p* < .001, η_p_^2^ = .30, a main effect of Response Number, *F*(2.44,109.58) = 909.07, *p* < .001, η_p_^2^ = .95, and an interaction between Drug and Response Number, *F*(3.68,165.62) = 4.38, *p* = .003, η_p_^2^ = .09. However, poorer accuracy was not associated with faster response times. Instead, response times were actually slower overall. A repeated measures ANVOA examining response times showed a main effect of Drug, *F*(1,45) = 21.68, *p* = .014, η_p_^2^ = .13, a main effect of Response Number, *F*(1.13,50.99) = 1026.79, *p* < .001, η_p_^2^ = .96, and a significant interaction between Response Number and Drug, *F*(1.13,50.64) = 11.61, *p* < .001, η_p_^2^ = .21.

### Effects of THC on mind wandering during the task (Exp 1)

In Exp 1 only, we used “thought probes” to assay the contents of participants’ thoughts while performing the working memory task (Fig 4A). THC significantly reduced reports of being On Task *t*(22) = 5.08, *p* < .001, *d* = 1.06. and increased frequency of both Mind Wandering *t*(22) = 4.42, *p* < .001, d = .92, and Zoning Out, *t*(22) = 2.13, *p* = .044, *d* = .45. In a separate ANOVA looking at the drug’s effects on type of mind wandering (Past, Future, Other, or “I Don’t Know”), there was no interaction between Drug and Mind Wandering Type, *F*(3,48) = 2.49, *p* = .07, η_p_^2^ = .135.

**Fig 4.**
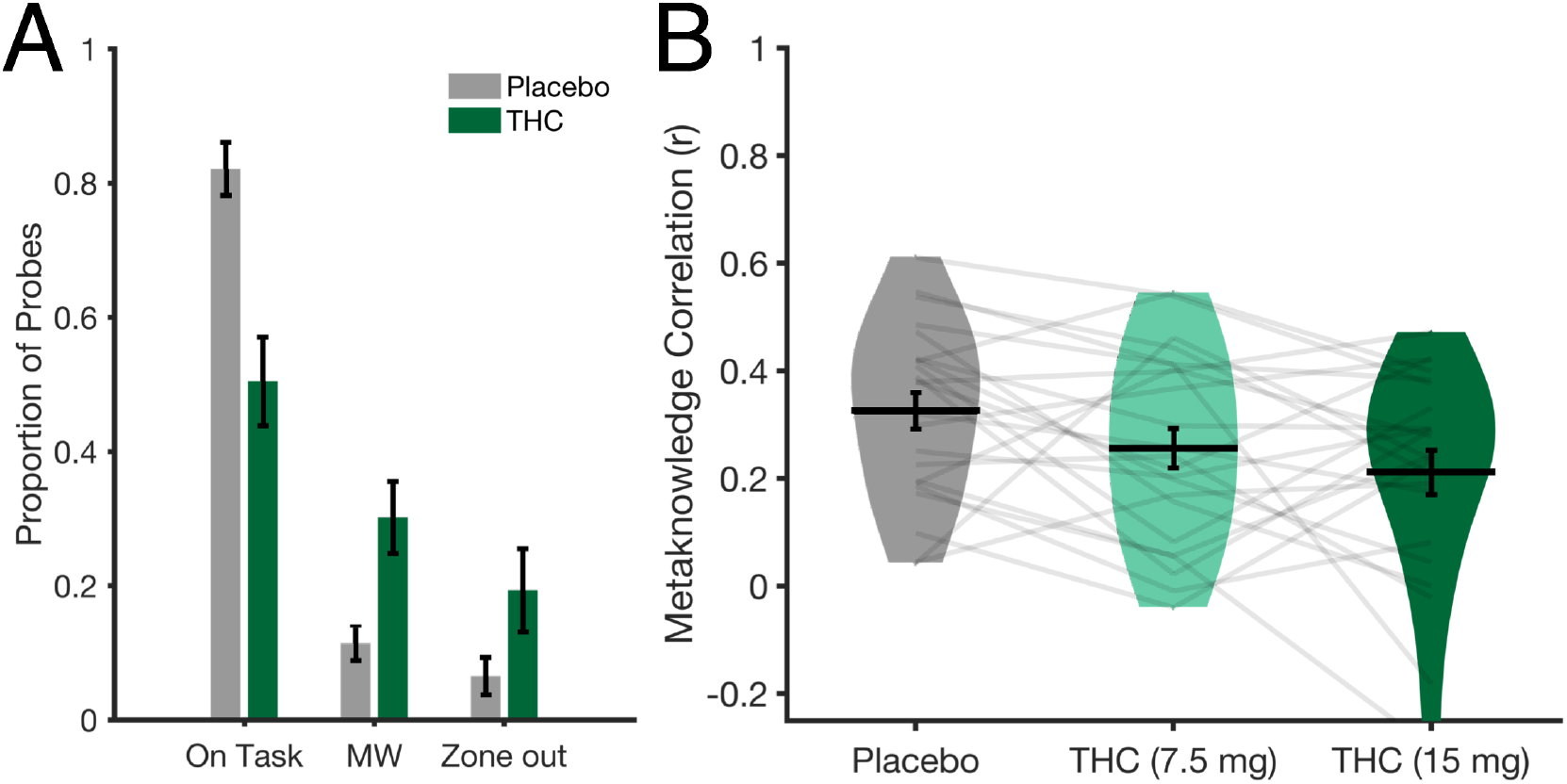
Changes to mind wandering and metacognitive accuracy after THC or placebo. (A) Mean changes to the distribution of thought probes in Exp 1 (placebo or 15 mg; n = 23). MW = Mind Wandering. Error bars represent 1 SEM. (B) Mean changes to metacognitive accuracy in Exp 2 (placebo, 7.5 and 15 mg; n = 23).

### Effect of THC on metacognitive accuracy (Exp 2)

In Exp 2 only, we examined subjects’ ability to accurately monitor ongoing task performance (i.e. metacognitive accuracy), by providing confidence ratings for each response. To measure metacognitive accuracy, we calculated the correlation between the number of *correct items* on each trial and the number of *confident responses* on each trial (separately for each individual using Spearman’s *r*). Higher, positive correlation values correspond with more accurate monitoring of task performance (e.g., the participant got N correct and reported N confident responses). We tested whether metacognitive accuracy declined as a function of Drug. A repeated measures ANOVA with factors Drug and Dose revealed a main effect of Drug, *F*(2,44) = 3.43, *p* = .04, η_p_^2^ = .14. Consistent with the impairment to overall performance, post-hoc *t*-tests for each dose revealed that this measure of metacognitive accuracy was impaired for the high dose (*p* = .03) but not the low dose (p = .07). We did not see an effect of the drug on metacognitive *bias* (mean number of confident items minus the mean number of correct items; positive numbers indicate over-confidence and negative numbers represent under-confidence). Participants were slightly overconfident in all three drug conditions (M = 0.51 items overconfident, SD = 0.97), and a repeated measures ANOVA revealed no significant main effect of drug condition on overconfidence, F(1.14,25.09) = 0.49, *p* = .52).

### Literature review and power analysis

Although the Digit Span task has been widely used, we found that the drug had no effect on this task in more than 70% (73.68%) of the 57 conditions that met the inclusion criteria of our Digit Span review (Fig 5A). A lack of sensitivity was evident for both Forward and Backward span, as well as other working memory tasks, such as Spatial N-Back (SI Results). The lack of effect was observed in studies using a range of doses including higher doses than what was used here, and multiple modes of administration (Table S1). The apparently weak effect of THC on working memory could indicate that the drug does not affect performance but, alternatively, it could reflect a lack of power in most prior studies. Although it was not possible to calculate effect sizes for the reviewed studies, we were able to demonstrate the effects of task time and sample size on power new empirical data.

**Fig 5.**
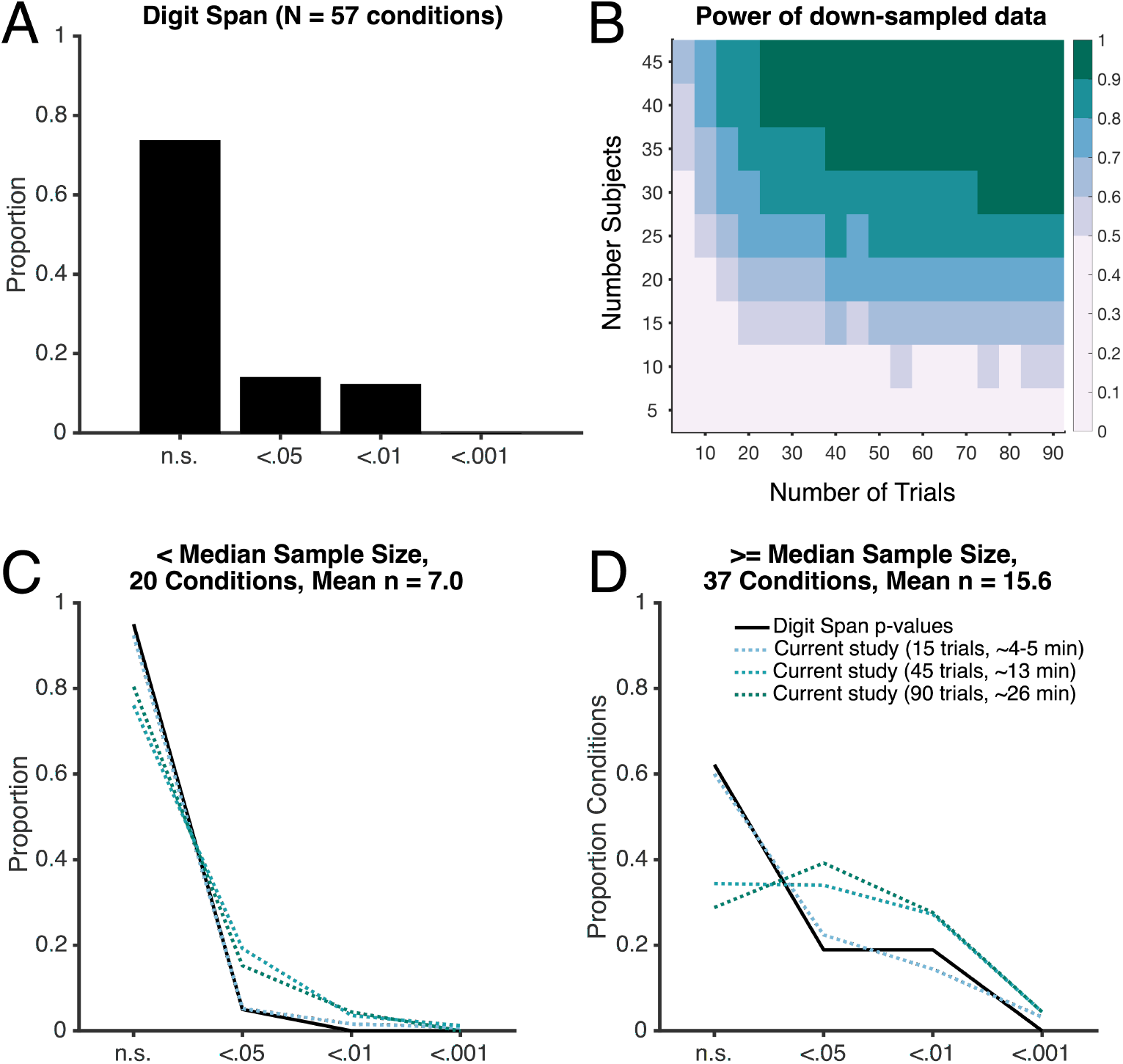
Power analysis predicts the distribution of *p*-values for published studies. (a) Histogram of reported *p*-values for 57 conditions of forward- and backward-digit span. (b) Down-sampling analysis of the current data. Each factorial combination of trial number (x-axis) and sample size (y-axis) contains the average power of 250 iterations sampled from the full data-set (collapsed across Exp 1 and the 15 mg of THC condition in Exp 2) (c-d) The black line shows the distribution of digit span *p*-values from the literature for conditions with fewer (c) or more (d) than the median number of subjects. Dotted lines show that this distribution of *p*-values from sampling the current visual working memory dataset with equivalent, insufficient power (e.g. 7 subjects and 15 trials)

To test whether the distribution of *p*-values in the literature review was related to insufficient statistical power or a lack of an effect of THC on working memory performance, we performed a down-sampling procedure [15] on the combined data from Exp 1 and Exp 2 (15 mg THC; n = 46, trials = 90). When we reduced the sample size and task length of our dataset to match those in the literature, we could nearly perfectly predict the distribution of p-values that was observed in the literature.

With 46 subjects and 90 trials per subject, the achieved power (1 - β) for our main effect of THC on working memory performance was in excess of 0.99. Fig 5B reveals the results of iterative down-sampling of this data (e.g. randomly choosing N subjects and T trials, calculating power). Each cell in this figure contains the average power for 250 random iterations. When down-sampling to a typical sample size and task duration for the Digit Span literature (e.g. 15 subjects, 15 trials), power plummets to only 0.47. In Figs 5C and 5D, we have plotted the distribution of *p*-values for Digit Span studies above and below the median sample size found in the literature. To compare predictions from our down-sampling procedure, we chose the cell from Fig 5B that most closely matched the number of subjects (5 subjects for the reviewed studies with below the median number of subjects overall, 15 subjects for the reviewed studies with above the median number of subjects overall), and then discretized the p-value outcomes for each of the 250 iterations (<.001, <.01, <.05, or n.s.). The digit span typically comprises 3 trials of each of 4-5 set sizes (12-15 trials) and takes approximately 4 – 5 minutes; 15 trials of the whole-report task likewise takes ∼5 min. As shown in Fig 5, with 5 minutes of task time and fewer subjects per “experiment”, we obtain distributions of *p*-values that are nearly identical to those in the empirical literature (Fig 5C, 5D).

## Discussion

The present study demonstrated that a single 15 mg dose of THC impairs working memory when tested under rigorous, placebo-controlled, double-blind conditions. THC, the main psychoactive constituent in cannabis, is commonly thought to impair working memory, but this effect has been difficult to demonstrate in controlled studies. We review previous studies assessing the effects of THC, and conclude that the failures to detect effects were quite likely due to inadequate statistical power. Using a well-powered sample (combined *n* = 46, [1-β] > .99), we found that a single 15 mg dose of THC reliably impairs visual working memory. This effect was not, however, apparent for a lower 7.5 mg dose (Experiment 2, n = 23).

We found a robust behavioral effect of THC on working memory performance, but more work is needed to understand the neural mechanisms underlying this behavioral deficit. For example, disruption to working memory is a key cognitive deficit in people with schizophrenia [77], and this deficit is hypothesized to be related to disruptions of the endocannabinoid system [78], more specifically to the dorsolateral prefrontal cortex (DLPFC)[79–81], but see [82]. The DLPFC is laden with CB1 receptors, the primary target of THC, and is a critical component of WM maintenance [83–85]. Studies of the neural mechanisms underlying WM disruption under the effects of THC may thus provide a reversible demonstration of WM deficits related to disruptions of the endocannabinoid system.

In addition to decreasing WM performance, subjective thought probes revealed that THC increased rates of mind wandering and zoning out (Exp 1) and decreased metacognitive accuracy (Exp 2). To our knowledge, our work provides the first demonstration of THC’s effects on mind wandering during a concurrent cognitive task. These finding are consistent with prior work on THC, including task-independent reports of mind wandering in structured interviews [86,87], failure to de-activate the default mode network during task performance [88] (but see [89]), and decreased error monitoring [42,43,90]. Similar to the effects of nicotine cravings [40] and alcohol [41], THC appears to increase mind wandering and other off-task mental states (e.g., “zoning out” or “mind blanking”[55]), and decrease awareness of task performance. These broad effects on conscious experience are likely to drive performance decrements in a broad range of cognitive tasks.

### Limitations and implications for future studies of THC and cognition

There in an intense public interest in the effects of cannabis on cognition and an urgent need for practical information that will guide use. Despite the importance of this study in providing new information on how THC affects memory, the study also had limitations that suggest future avenues for research. First, we were able to test working memory at only one, relatively late time-point after oral consumption of a THC capsule (160 – 220 minutes), and at two moderate doses [91]. It will be important to characterize the effects of THC on working memory performance over the full time-course of the drug, and, importantly at higher doses and by different routes of administration (especially smoked and vaped). In addition, further work is needed to understand relationships between THC-related working memory impairments and other task and participant factors, such as recent and lifetime exposure to THC, and generalizability to other cognitive constructs. Second, we need more information on the severity of working memory disruption and the extent to which the effect depends on initial performance. Although the effect we observed was relatively large (*d* = .65), this effect is smaller than, for example, normal variation in working memory performance across individuals. The behavioral performance difference between the placebo and drug conditions was 0.29 items, but the difference between the top- and bottom-half of individuals within the placebo condition was 0.80 items. Further, the effects of the drug may be especially pronounced in certain at-risk populations, including those with initially poor working memory performance.

Our finding that power curves for a down-sampled visual working memory task closely matched the observed power in the Digit Span literature suggests that task length is a strong driving factor in the low rate of positive drug effects in the literature. However, more methodological work is needed to determine if directly manipulating the task length of standard tasks (e.g., a 25-minute rather than 5-minute version of the Digit Span task) will rescue statistical power as our simulations suggest. Alternatively, it is possible that other task differences (e.g., factor loadings) also contribute to the low rate of positive drug effects in the literature. For example, simple span tasks (e.g., the Forward Digit Span) do not load well onto a general working memory factor at the latent level [20], and often fail to predict individual differences in general fluid intelligence [19,92–95], but also see [96]. Other common tasks (e.g., spatial *N*-back, Table S2) likewise have potential differences from the specific visual working memory task used here. N-back tasks load well onto a general WM factor at the latent level [20] and are useful for investigating the executive function component of WM (vs. the storage component). However, *N*-back tasks are not highly correlated with other working memory tasks [97–99] and often have relatively poor statistical reliability [95], potentially making it difficult to detect effects across treatment conditions.

Our literature search revealed the importance of statistical power in studies of THC on cognition. Here, we used one relatively long task (∼30 minutes, 90 trials). In contrast, many earlier studies used several shorter tasks (e.g. ten 3-minute tasks), presumably to assess a range of potential deficits. However, because task length and statistical power have direct tradeoffs, this approach may miss important effects. Similar problems of inadequate power may exist in other studies of effects of drugs on working memory and other aspects of cognition. Thus, we recommend that longer tasks be used to determine the effects of drugs on cognition.

## Contributions

Manoj Doss and Elisa Pabon collected data. Kirsten Adam performed analyses and drafted the manuscript. Kirsten Adam, Manoj Doss, Elisa Pabon, Edward Vogel, and Harriet de Wit planned the experiments and revised the manuscript.

## Footnotes

* Specific to the Discrete Whole Report task, previously reported reliabilities include: Cronbach’s alpha for stability of performance across blocks > .9 [17], split-half reliability, with training sessions over multiple weeks, *r* = .60 - .86 [18], current study split-half reliability *r* >= .90).

† Greenhouse-Geisser corrected values are reported whenever the assumption of sphericity is violated.

