## Supplemental Information for "Δ^9^-Tetrahydrocannabinol (THC) impairs visual working memory performance: A randomized crossover trial"

### Supporting Information

#### SI Methods

##### CONSORT information for study protocols

**Registration information.** Study 1 was part of study, Effects of THC on Emotional Memory Retrieval (TARE), Clinical Trials ID NCT03471585, protocol available at: https://clinicaltrials.gov/ct2/show/NCT03471585. Study 2 was part of study, "Developing a Mobile Method to Measure THC-induced Impairment (AIS)", Clinical Trials ID NCT03804840, protocol available at: https://clinicaltrials.gov/ct2/show/record/NCT03804840. Study recruitment and data collection occurred during the periods February 2017 to October 2017 (Study 1) and between March 2017 to December 2018 (Study 2). Study recruitment was ended when the *a priori* sample size was reached. Recruitment and attrition information can be found in Figures S3 and S4.

**Outcomes. Study 1:** Primary outcomes for this study were the pre-registered physiological effects, subjective effects, and working memory performance. The mind-wandering measure during the working memory test was a secondary measure added after the clinical trials registration was uploaded but prior to any data were collected. **Study 2:** Primary outcomes for this study were "performance on basic cognitive and psychomotor tasks as well as perceived performance. Outcomes reported here (working memory task performance, perceived number of items correct, standard physiological and subjective effects) were finalized after the clinical trial registration was uploaded, but before any data were collected.

**Important harms and unintended effects.** No important harms or unintended effects were noted in Study 1. In Study 2, three participants reported feeling anxious in the high dose (15 mg THC) condition.

##### Subjective measures

**Addition Research Center Inventory (ARCI).** The Addiction Research Center Inventory (ARCI) is set of 53 true/false questions to measure drug-specific behavioral and mood effects. These questions include the 49 questions reported in Martin et al. [1] and an additional 4 questions from the Marijuana group sub-scale. The 53-item ARCI measure contains a sub-set of items from the following scales reported by Haertzen [2]: The Pentobarbital, Chlorpromazine, & Alcohol Group Scale (PCAG; sedation); The Morphine-Benzedrine Group Scale (MBG; euphoria); the Lysergic Acid Diethlamide Group Scale (LSD; dysphoric /psychotomimetic); the Benzedrine Group Scale (BG; empiric amphetamine scale); Marijuana Group Scale (Ma). In addition, some items in the set of 53 were devised by Martin et al. [1] as part of a revised amphetamine scale. From this set of 53 questions, we focused our analyses on the 12-item "revised Marijuana scale" [3] as this scale is tailored to the effects of cannabis.

**Drug Effects Questionnaire (DEQ).** The Drug Effects Questionnaire (DEQ) is a set of 5 questions about the current effects of a drug [4]. For each question, participants rate how much they feel a drug effect, like the effect, dislike the effect, feel high, and want more of the drug. Participants rate each question on a continuous sliding scale from “not at all/neutral” to “very much.” Participants were instructed to select “not at all/neutral” if they had not yet received a capsule (first time-point).

**Visual Analog Scales (VAS).** The Visual Analog Scales (VAS) are measures used to assess individual dimensions of subjective mood. These measures were only used in Exp 1. We used a version of this scale with 13 adjectives [5] – anxious, stimulated, sedated, elated, insightful, sociable, confident, lonely, playful, dizzy, loving, friendly, and restless. Participants rated the degree to which they felt each of these adjectives on a continuous sliding scale from “not at all” to “extremely.”

**End of Session Questionnaire.** The end of session questionnaire (ESQ) asked participants which class of drugs they thought they had received during the current session.

**Design**

**Time points for cardiovascular and subjective measures.**

*Experiment 1:* Cardiovascular measures were taken at baseline (immediately preceding capsule administration) and at 30, 60, 90, 120, 190 and 210 min post-capsule. Subjective measures were taken at the same time-points as the cardiovascular measures, with the exception of the 210 min time point. To assess drug effects relative to baseline, we used the time-point closest in time to the WM test (120 min for 4 participants who started the WM task at ~130 min; 190 min for remaining participants who started the WM test at 160 min).

*Experiment 2:* Cardiovascular and subjective measures were taken at baseline and at 30, 60, 90, 150 and 240 min post-capsule. To calculate change from baseline, we used the time-point immediately following the WM Test (240 min).

**Additional design details for Exp 1.** The working memory task stimuli (“Discrete Whole-Report Task”) are depicted in Figure 1 in the main text. In Exp 1, participants performed the working memory task in two total sessions, but these two sessions were part of a larger, multi-session study, as reported in Doss et al. [6]. In this design, participants came into the lab for one orientation pre-session and then a total of six experimental sessions. This multi-session design was used in order to disentangle the effects of THC on long-term memory encoding versus retrieval. To this end, participants performed both a long-term memory retrieval test and a long-term memory encoding test during capsule administration. During Session 1, participants encoded two different sets of stimuli (emotional stimuli; Deese-Roediger-McDermott stimuli). After 48 hours, they returned for Session 2. During Session 2, they took a capsule (THC or placebo), were tested on their memory of the stimuli they had learned during Session 1, encoded a new set of stimuli (object/scene) into long-term memory, and completed the working memory test. Finally, after 48 hours, the participants returned for Session 3 and were tested on their memory of the object/scene stimuli that were learned in Session 2. After a minimum of 72 hours, participants returned for Session 4. Sessions 4-6 followed the exact procedures as Sessions 1-3, with the exception that the specific stimuli used for each long-term memory test were unique (i.e., different images). This six-session design allowed for a within-subjects comparison of the encoding- and retrieval-specific effects of THC on long-term memory performance.

**Procedures**

**Time of abstinence.** Participants were instructed that each session would begin with a drug test. In Exp 1, participants were instructed to abstain from using alcohol, prescription drugs (except contraceptives), and over-the-counter drugs for 24 hours prior to the encoding sessions (e.g. Session 1) and to remain abstinent through the corresponding retrieval session (e.g. Session 3). They were instructed to abstain from cannabis starting 1 week before the encoding sessions, and other elicits drugs starting 48 hours before the encoding sessions. In Exp 2, participants were instructed to abstain from the above-named substances for at least 48 hours before each experimental session; participants were also cautioned that a negative drug test would be required to participate in the study, and some substances may take longer than 48 hours to leave the system.

**Exp 1.** Participants completed other cognitive tasks before completing the working memory task in Exp 1. Cognitive testing began at approximately 120 minutes after capsule consumption. First, participants were tested on their memory for emotional stimuli that had been encoded 48 hours prior (30 min). Second, participants were tested on their memory for the DRM stimuli (5 min). Third, participants encoded a new set of stimuli (object/scene; 20 min). Finally, participants completed the working memory task that is the central focus of this paper.

**Exp 2**. Participants also completed other cognitive tasks before completing the working memory task in Exp 2; cognitive testing likewise began at ~120 minutes after capsule consumption. Participants completed a series of tasks on a desktop computer and another series of tasks on a mobile phone app. The order of the desktop versus app tasks was counter-balanced. The desktop tasks took approximately 30 min, and the phone tasks took approximately 5 min. After completing both the desktop and phone tasks, participants were allowed to relax for 30 min. Then, they completed both sets of tasks a second time. Finally, participants performed the working memory task (~220 min post-capsule).

**Literature search**

We examined the literature to gauge the strength of prior evidence that THC impairs working memory. We searched for all studies using either a Digit Span task, Digit Recall Task, or Spatial *n*-back Task (Tables S1 and S2). In addition, we also report additional recent within-subjects, placebo-controlled studies that used visual working memory tasks and some miscellaneous tasks with a working memory component (Table S3). Inclusion criteria were (1) human subjects, (2) testing Δ9-tetrahydrocannabinol, either cannabis (alone, e.g. not mixed with tobacco containing nicotine) or synthetically produced THC, e.g. Marinol® or Dronabinol, (3) placebo-controlled, (4) within-subjects design with randomized order of administration, and (5) report running statistical tests.

We performed our initial literature search in July 2018, and updated it in August 2019. We did not formally exclude papers based on publication date, but we did not find any studies meeting our inclusion criteria from before 1970. We began our search for papers based on previous literature reviews that have discussed the acute effects of THC on working memory [7–12]. Importantly, these reviews included two systematic reviews of the effects of THC on behavioral performance. Both papers included a section on the acute effects of THC on working memory performance. Zuurman and colleagues [12] systematically reviewed the literature up until 15 Nov 2007. Broyd and colleagues [11] systematically reviewed the literature from 2004 until February 2015. To supplement these 2 systematic reviews, we performed a mini-systematic review of studies published from 1 Jan 2015 – 29 August 2019. In this mini-review, we searched Scopus for articles (English-language only) containing a word related to a THC manipulation ("cannabis" or "tetrahydrocannabinol", or "marijuana" or "marihuana") and a working memory task ("digit span", "digit recall", "working memory", "n-back", "short-term memory", or "immediate memory"). To reduce the number of results, we excluded the exact keywords "Nonhuman" and "Animals".

Beginning with the cited systematic and qualitative reviews, we examined papers cited as containing a working memory task and then looked at the forward- and backwards-citations of these papers for other relevant papers. Working memory task results were often part of a larger test battery and often yielded a null result, so these measures were often not reported in the abstract and reported in a single line (e.g. "no other measures showed significant effects"). We focused, in particular, on finding all studies using the Digit Span task and the Spatial N-back task, which were overwhelmingly the most commonly used tasks. We also include a summary of studies using other commonly used tasks (particularly, visual working memory tasks), but chose not to include some older tasks (e.g., Serial Sevens).

Some studies have been previously discussed in review papers but did not meet our inclusion criteria. For example, Heishman et al. [13] has typically been included in reviews on the acute effects of THC on working memory / cognitive performance [7,8], but this paper does not report any group-level statistics for the working memory task. They reported that "2 out of 3 subjects showed impairment" but did not quantify this impairment formally at the individual or the group level. Thus, this paper did not meet the inclusion criteria for our literature review. Likewise, other studies cited in some previous reviews did not control for the effects of time on task performance (i.e. they did not counterbalance the administration of THC and placebo.

Many studies that administered THC via cigarette reported the percentage of THC contained in 1 cigarette. To make comparisons across studies easier, we have included the estimated mg of THC in the cigarette that was smoked by participants based on an estimated cigarette weight of 800 mg, based on published work on the weight of NIDA-supplied cigarettes [14]. Note, in some studies participants consumed THC as a function of the number of "puffs". For these studies we cannot report administered THC in mg, as the exact conversion is unknown. We have chosen to include studies in which THC was ingested, smoked, or administered intravenously. In addition, some THC was synthetic (e.g. Dronabinol or Marinol®) and some was "natural" (e.g. cannabis plant). Previous work has shown that the subjective and physiological effects are comparable for synthetic and natural THC [15]. The time course is much slower for ingested THC relative to smoked THC for a comparable dose (e.g. ~8 mg), but the subjective and physiological effects at the peak response are comparable [15]. Conversely, a smaller intravenous dose yields stronger physiological and subjective responses relative to an equivalent weight of smoked THC (e.g. 5 mg intravenous vs. 14 mg smoked), but the time course of intravenous and smoked THC is roughly equivalent [16].

#### SI Results

**Cannabis experience did not modulate the drug effect.**

We found no noticeable relationship between a subject's previous experience with THC and overall impairment of working memory by THC in these studies. As noted in the main body of the text, there was no significant difference in the drug effect across experiments ($p>.05$), justifying the combination of experiments for the across-experiment analyses. Yet, there was a large difference in the average lifetime number of uses of cannabis ($p < .008$) and monthly uses of cannabis ($p < .02$) for the two experiments because of differences in recruitment criteria across the two studies. Thus, experience with THC cannot explain the impairment of WMC by working memory performance in the current set of experiments.

**Correct identification of administered drug did not modulate the drug effect.**

We found no evidence to support the idea that participants who correctly guessed which drug they were administered had a smaller or larger sized drug effect compared to participants who did not correctly guess. Participants in our studies were not informed in advance that they would receive either a cannabinoid or a placebo; they were informed that they might receive one of many possible drugs (cannabinoid, sedative, stimulant) or a placebo. In the end of session questionnaire, they guessed which drug (or placebo) they had received during the session (Table S4).

In Experiment 1, the overall working memory effect was not significantly different when comparing effects (Mean number correct for THC – Placebo) of participants who identified both the placebo and drug sessions correctly (n = 12, Mean difference = -.32 items, SD = .55) versus those who got at least one session incorrect (n = 11, Mean difference = -.34 items, SD = .29), $t(21) = 0.07$, $p = .94$. Similarly there was no difference between those participants who correctly identified the drug session as a cannabinoid (n = 17, Mean difference = -.34, SD = .48) with those who did not correctly identify the drug session as a cannabinoid ($n = 6$, Mean difference = -.32, SD .28), $t(21) = 0.13$, $p = .89$.

In Experiment 2, the overall working memory effect (15 mg drug session versus Placebo) was similarly not significantly different when comparing the mean drug effect for participants who identified both the placebo and drug sessions correctly (n = 12, Mean difference = -.16 items, SD = .46) versus those who got at least one session incorrect (n = 11, Mean difference = -.33 items, SD = .45), $t(21) = 0.91$, $p = .37$. Similarly there was no difference between those participants who correctly identified the drug session as a cannabinoid (n = 15, Mean difference = -.25, SD = .56) with those who did not correctly identify the drug session as a cannabinoid ($n = 8$, Mean difference = -.22, SD .14), $t(21) = 0.11$, $p = .91$.

**Effects of sex, body mass index, physical weight, and physical height mass index on working memory impairment by THC**

In the combined sample (Exps. 1 and 2, n = 46), we looked at potential modulations of the working memory impairment effect by reported biological sex, physical weight, physical height, and body mass. We found a small but significant effect of sex (Figure 1), with women showing more severe working memory impairment, mean difference = 0.25 items, $t(44) = 2.05$, $p = .047$. However, this small difference does not fit with prior work reporting no sex difference in THC kinetics and metabolite composition [17]. We did find a significant correlation between BMI and working memory impairment: those with a larger BMI showed *more* severe WM impairment, $r = -.35$, $p = .017$. For ease of comparison to the sex effect, we conducted a median split and found a robust difference in WM impairment for those with high and low BMI, $t(44) = 3.50$, $p = .001$, but no average difference between men and women in BMI, $t(44) = 0.84$, $p = .41$. The prolonged effect in individuals with higher BMI is consistent with the fact that THC is lipophilic: participants with a higher percentage of body fat may show prolonged effects of THC intoxication as the drug is sequestered in fat.

Likewise, despite expected sex differences in both height ($p = .017$) and weight ($p < 1 \times 10^{-3}$), there was no significant correlation between working memory impairment and either height (p = .71) or weight (p = .38). The median split analyses of height and weight were likewise null effects (height, p = .94; weight, p = .36). This preliminary effect suggests that individuals with a high body mass index (and/or body fat percentage) may show prolonged effects of THC intoxication, though further work is needed on this topic.

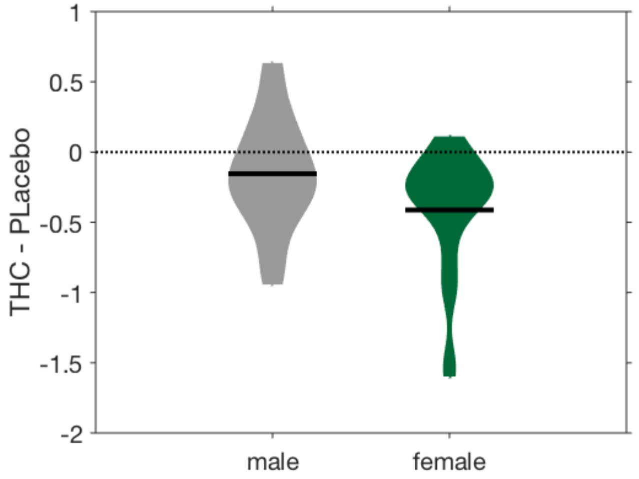

**Figure S1. Working memory impairment by sex.** There was a small but significant modulation of the working memory impairment effect by sex ($p = .047$).

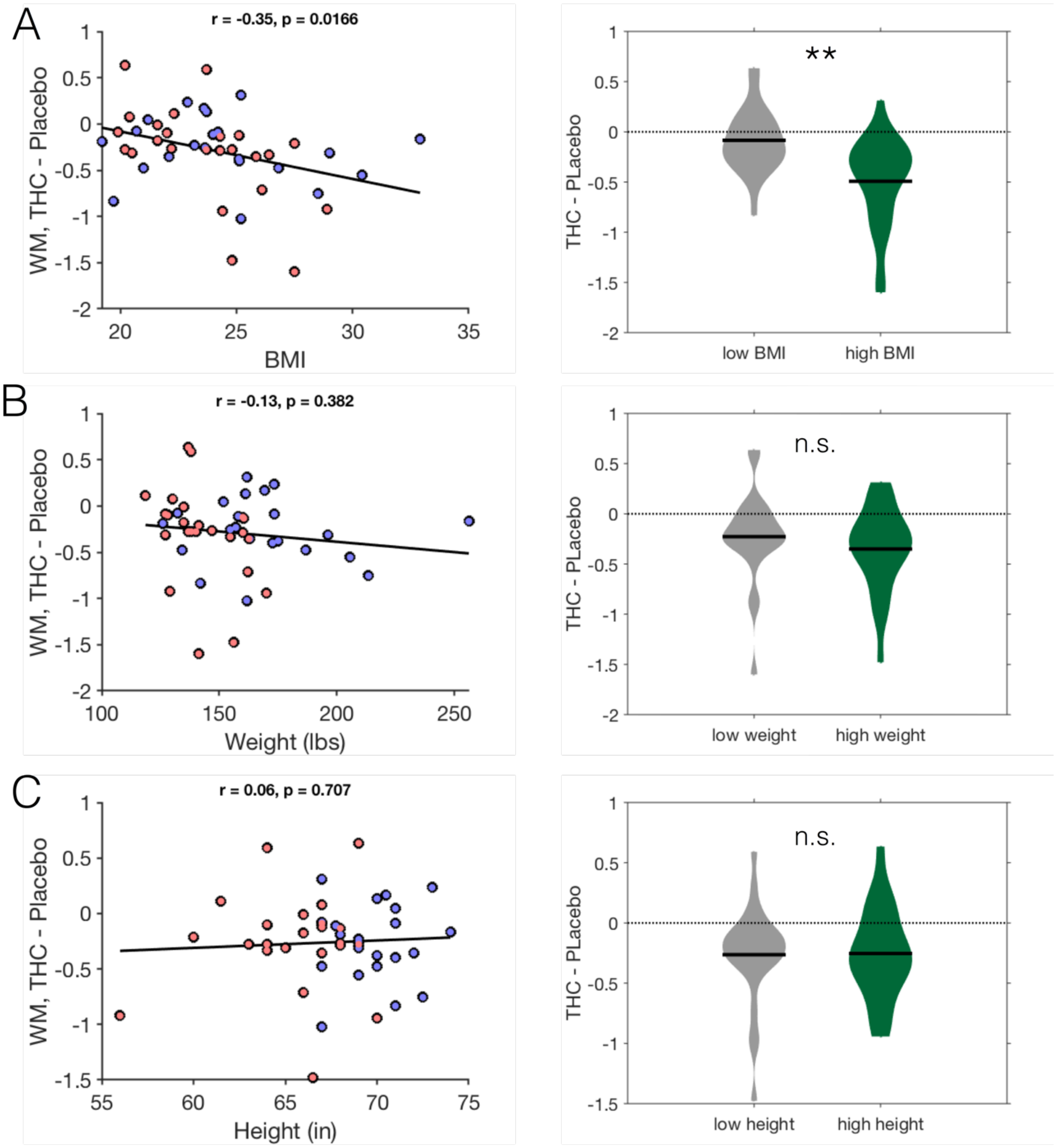

**Figure S2. Working memory impairment by body mass index (BMI), weight, and height.** (A) Body mass index was significantly correlated with working memory impairment. Those with a higher body mass index showed more impairment. In the correlation plots, blue dots represent men and red dots represent women (for illustrative purposes only). The right panel of A shows a median split analysis. (B) There was no correlation between physical weight and working memory impairment. (C) There was no correlation between physical height and working memory impairment.

**Figure S3. CONSORT flow-chart for Study 1.** This chart has been adapted from the standard 2-group format to one more appropriate for a within-subjects crossover design (see Dwan et al. 2019). P = placebo, D = drug (15 mg THC).

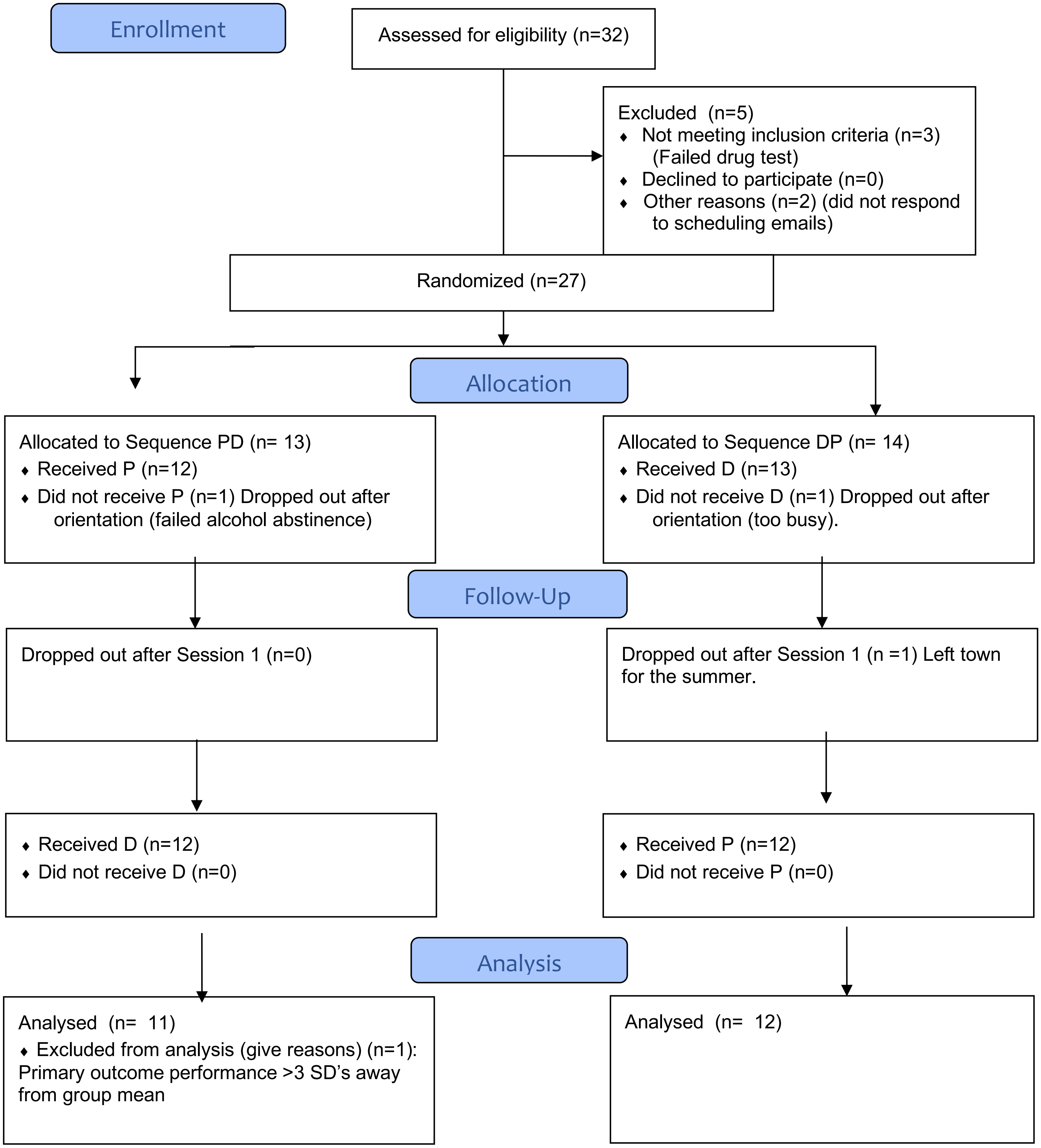

**Figure S4. CONSORT flow-chart for Study 2.** This chart has been adapted from the standard 2-group format to one more appropriate for a within-subjects crossover design (see Dwan et al. 2019). P = placebo, L = low dose (7.5 mg), H = high dose (15 mg).

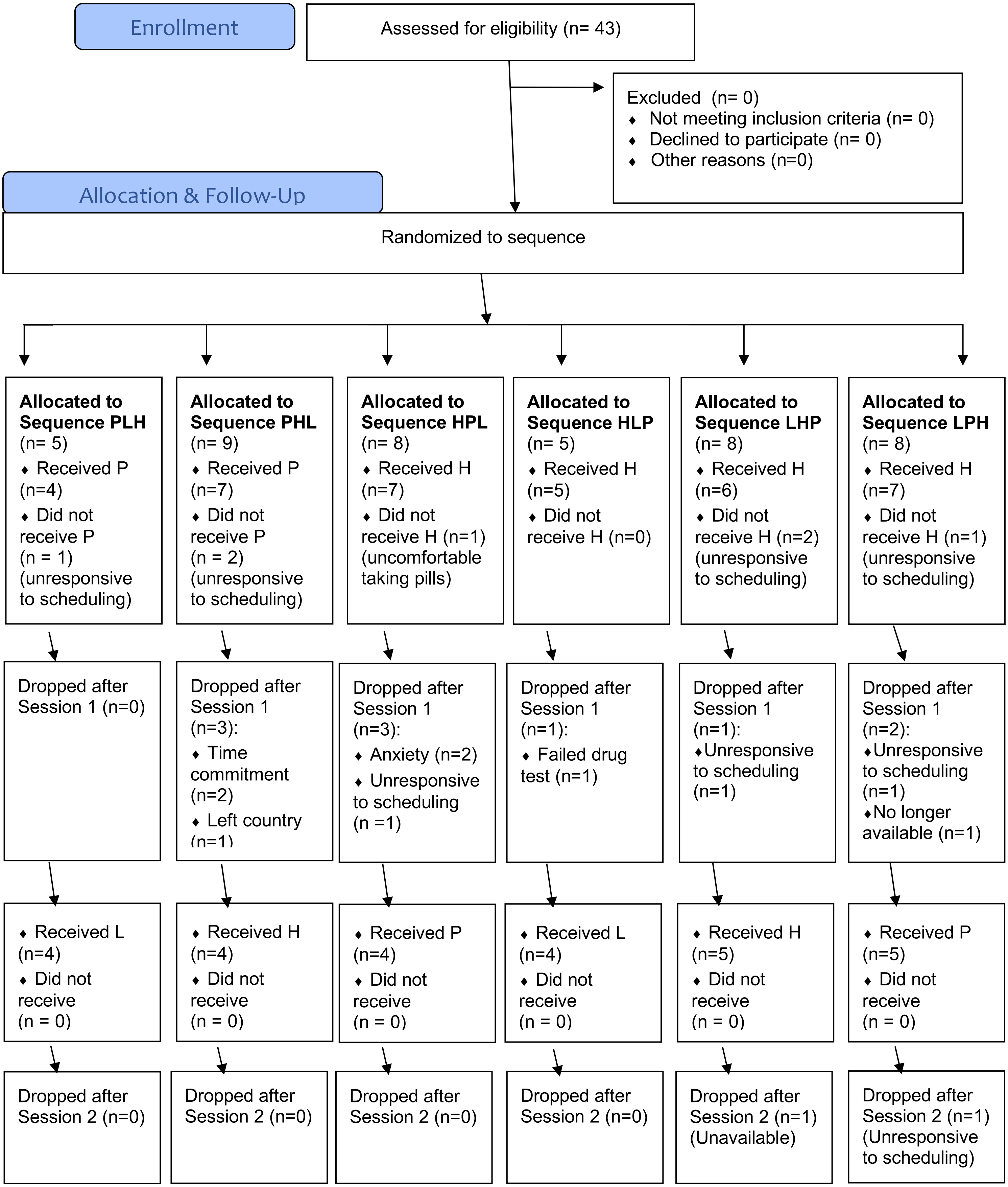

| | | | | | |
|---|---|---|---|---|---|
| ♦ Received H (n=4)<br>♦ Did not receive (n = 0) | ♦ Received L (n=4)<br>♦ Did not receive (n = 0) | ♦ Received L (n=4)<br>♦ Did not receive (n = 0) | ♦ Received P (n=4)<br>♦ Did not receive (n = 0) | ♦ Received P (n=4)<br>♦ Did not receive (n = 0) | ♦ Received H (n=4)<br>♦ Did not receive (n = 0) |

**Analysis**

| | | | | | |
|---|---|---|---|---|---|
| ♦ Analyzed WM (n=4)<br>♦ Excluded from analysis (n=0) | ♦ Analyzed WM (n=3)<br>♦ Excluded from analysis (n=1) (WM performance > 3 SD's from group average) | ♦ Analyzed WM (n=4)<br>♦ Excluded from analysis (n=0) | ♦ Analyzed WM (n=4)<br>♦ Excluded from analysis (n=0) | ♦ Analyzed WM (n=4)<br>♦ Excluded from analysis (n=0) | ♦ Analyzed WM (n=4)<br>♦ Excluded from analysis (n=0) |

#### Literature search

Conditions that were reported separately appear as separate lines in the tables below. For example, most studies separately reported the statistics for forward span versus backward span, and most studies separately reported the statistics for the different doses administered to participants. Because of inconsistent reporting standards across studies, it was not possible to recalculate effects when collapsed across all conditions or to calculate effect sizes for each condition. Where possible, we present exact *p*-values. However, *p*-values were rounded to $p < .01$, $p < .05$, and “not significant” (n.s.) in the vast majority of studies.

**Digit span and recall tasks.** We found a total of 21 papers that reported a measure of digit span or digit recall and met our inclusion criteria (Table S1). Of these, 15 included a Forward Digit Span Task or a Backward Digit Span Task [18–32]. These 15 studies contained a total of 57 conditions (e.g. different drug doses).

**Table S1.** Literature summary of within-subjects, placebo-controlled studies testing Working Memory with a Digit Span (“Span”) or Digit Recall (“Recall”) Task.

| Study | Ref. | n | Route | Task | Condition - Task | Condition - Dose | *p*-value |
|---|---|---|---|---|---|---|---|
| Tinklenberg et al. 1970 | [18] | 8 | Oral | Span | Forward | 20 mg | < .05 |
| | | | | | | 40 mg | < .05 |
| | | | | | | 60 mg | < .05 |
| | | | | | Backward | 20 mg | <.01 |
| | | | | | | 40 mg | <.01 |
| | | | | | | 60 mg | <.01 |
| Melges et al. 1970 | [19] | 8 | Oral | Span | Forward | Mean: 20, 40, 60 mg | 0.05 |
| | | | | | Backward | Mean: 20, 40, 60 mg | 0.01 |
| Tinklenberg et al. 1972 | [33] | 15 | Oral | Recall | Forward (10 - 20 digits) | 26 mg | n.s. |
| Dornbush & Kokkevi (1976) | [32] | 20 | Cigarette | Span | Forward | 78 mg cannabis leaf, percentage THC not reported (30 minutes) | n.s. |
| | | | | | | 78 mg cannabis leaf, percentage THC not reported (70 minutes) | n.s. |
| | | | | | Backward | 78 mg cannabis leaf, percentage THC not reported (30 minutes) | n.s. |
| | | | | | | 78 mg cannabis leaf, percentage THC not reported (70 minutes) | n.s. |
| McDonald et al. 2003 | [20] | 37 | Oral | Span | Forward | 7.5 mg | n.s. |
| | | | | | | 15 mg | n.s. |
| | | | | | Backward | 7.5 mg | <.05 |
| | | | | | | 15 mg | <.05 |
| Ballard & de Wit 2011 | [21] | 11 | Oral | Span | Forward | 2.5 mg | n.s. |
| | | | | | Backward | 2.5 mg | n.s. |
| Casswell et al. 1973 | [22] | 12 | Cigarette | Span | Forward | 3.3. mg | n.s. |
| | | | | | | 6.6 mg | n.s. |
| | | | | | Backward | 3.3 mg | n.s. |
| | | | | | | 6.6 mg | n.s. |

| | | | | | | | |
|---|---|---|---|---|---|---|---|
| Galanter et al. 1973 | [34] | 12 | Cigarette | Recall | Forward (9 digits) | 10 mg "natural" | p<.01 |
| | | | | | | 10 mg synthetic | p<.05 |
| | | | | | Backward (9 digits) | 10 mg "natural" | p<.01 |
| | | | | | | 10 mg synthetic | p<.01 |
| Cappell & Pliner 1973 | [35] | 20 | Cigarette | Recall | Backward (9 digits) | 12 mg | n.s. |
| Hooker & Jones 1987 | [23] | 12 | Cigarette | Span | Forward | 10.7 mg | n.s. |
| | | | | | Backward | 10.7 mg THC | n.s. |
| Chait et al. 1988 | [24] | 8 | Cigarette | Span | Forward | 0 to 8 cumulative puffs, 1.4% THC | p<.01 |
| Heishman et al. 1989 | [25] | 12 | Cigarette | Span | Forward | 1.3% THC (~10 mg) | p<.01 |
| | | | | | | 2.7% THC (~22 mg) | n.s. |
| | | | | | Backward | 1.3% THC (~10 mg) | n.s. |
| | | | | | | 2.7% THC (~22 mg) | p<.01 |
| Zacny & Chait 1991 | [26] | 10 | Cigarette | Span | Backward | 4 puffs 2.3% THC (0 or 20 sec Breathhold Duration) | n.s. |
| Azorlosa et al. 1992 | [27] | 7 | Cigarette | Span | Forward | 4 puffs 1.75 % THC | n.s. |
| | | | | | | 10 puffs 1.75% THC | n.s. |
| | | | | | | 25 puffs 1.75% THC | n.s. |
| | | | | | | 4 puffs 3.55% THC | n.s. |
| | | | | | | 10 puffs 3.55% THC | n.s. |
| | | | | | | 25 puffs 3.55% THC | <.05 |
| | | | | | Backward | 4 puffs 1.75 % THC | n.s. |
| | | | | | | 10 puffs 1.75% THC | n.s. |
| | | | | | | 25 puffs 1.75% THC | n.s. |
| | | | | | | 4 puffs 3.55% THC | n.s. |
| | | | | | | 10 puffs 3.55% THC | n.s. |
| | | | | | | 25 puffs 3.55% THC | n.s. |
| Chait & Perry 1994 | [28] | 14 | Cigarette | Span | Backward | 3.6% THC (~29 mg) | n.s. |
| Azorlosa et al. 1995 | [29] | 7 | Cigarette | Span | Forward | 10 puffs 1.75% THC, 30 - 90 ml puff volume | n.s. |
| | | | | | | 10 puffs, 3.55% THC, 30 -90 mL puff volume | n.s. |
| | | | | Span | | 10 puffs 1.75% THC, 0 - 20 sec Breathhold Duration | n.s. |
| | | | | | | 10 puffs, 3.55% THC, 0 - 20 sec Breathhold Duration | n.s. |
| | | | | Span | Backward | 10 puffs 1.75% THC, 30 - 90 ml puff volume | n.s. |
| | | | | | | 10 puffs, 3.55% THC, 30 -90 mL puff volume | n.s. |
| | | | | Span | | 10 puffs 1.75% THC, 0 - 20 sec Breathhold Duration | n.s. |
| | | | | | | 10 puffs, 3.55% THC, 0 - 20 sec Breathhold Duration | n.s. |
| Fant et al. 1998 | [36] | 10 | Cigarette | Recall | Recall missing digit (9 digits) | 1.8% THC (~14.4 mg) | n.s. |
| | | | | | | 3.6% THC (~29 mg) | n.s. |
| Greenwald & Stitzer 2000 | [37] | 5 | Cigarette | Span | Combined | 18 puffs, 3.6% THC | 0.08 |
| Hart et al. 2001 | [30] | 18 | Cigarette | Span | Forward | 1.8% THC (~14.4 mg) | n.s. |
| | | | | | | 3.6% THC (~29 mg) | n.s. |

| | | | | | | | |
|---|---|---|---|---|---|---|---|
| | | | | | Backward | 1.8% THC (~14.4 mg) | n.s. |
| | | | | | | 3.6% THC (~29 mg) | n.s. |
| | | | | Recall | 1 trial of 8 digit recall | 1.8% THC (~14.4 mg) | n.s. |
| | | | | | | 3.6% THC (~29 mg) | <.05 |
| Ramesh et al. 2013 | [38] | 18 | Cigarette | Recall | 1 trial of 8 digit recall | 2 puffs 5.5%-6.2% THC | n.s. |
| | | | | | | 4 puffs 5.5%-6.2% THC | n.s. |
| | | | | | | 6 puffs 5.5%-6.2% THC | n.s. |
| Morrison et al. 2009 | [31] | 19 | Intravenous | Span | Forward | 2.5 mg | p<.05 |
| | | | | | Backward | 2.5 mg | p<.005 |

**Spatial n-back task.** We found a total of 9 papers that reported the results of a spatial n-back task and met our inclusion criteria (Table S2).

**Table S2.** Literature summary of within-subjects, placebo-controlled studies testing Working Memory with a Spatial *n*-back Task.

| Study | Year | n | Route | Dose | Condition - Task | p-value |
|---|---|---|---|---|---|---|
| Ilan et al. 2004 | [39] | 10 | Cigarette | 3.5% THC (~28 mg) | 0-back | n.s. |
| | | | | | 2-back | <.05 |
| Ilan et al. 2005 | [40] | 23 | Cigarette | Mean: 1.8% / 3.6% THC (~14/29 mg) | 1-back, 0:20 min | <.05 |
| | | | | | 1-back, 1:20 min | n.s. |
| | | | | | 1-back, 2:20 min | n.s. |
| | | | | | 2-back, 0:20 min | <.05 |
| | | | | | 2-back, 1:20 min | <.05 |
| | | | | | 2-back, 2:20 min | <.05 |
| Hart et al. 2010 | [41] | 24 | Cigarette | 1.8% THC (~14.4 mg) | 1-back | n.s. |
| | | | | | 2-back | n.s. |
| | | | | 3.9% THC (~31 mg) | 1-back | n.s. |
| | | | | | 2-back | n.s. |
| Dumont et al. 2011 | [42] | 16 | Vapor | 4 + 6 + 6 mg (90 min intervals) | 1-back | <.01 |
| | | | | | 2-back | n.s. |
| | | | | | 3-back | n.s. |
| Morrison et al. 2011 | [43] | 16 | Intravenous | 1.25 mg | Main effect: 0-, 1- and 2- back | .11 |
| Desrosiers et al. 2015 | [44] | 25 | Cigarette | 6.8% THC (54 mg) | 1-back | n.s. |
| | | | | | 2-back | <.05 |
| | | | | | 3-back | n.s. |
| Mokrysz et al. 2016 | [45] | 40 | Vapor | .1068 mg / kg (M = ~7.5 mg) | Main effect: 0-, 1- and 2- back | <.001 |
| Hindocha et al. 2017 | [46] | 24 | Cigarette | 66.7 mg Bedrobinol (10.7 mg THC) | 0-back | n.s. |
| | | | | | 1-back | .018 |
| | | | | | 2-back | <.001 |
| Morgan et al. 2018 | [47] | 48 | Vapor | 8 mg THC | Main effect: 1- and 2- back | .012 |

**Other tasks.** In Table S3 we report other papers that we include a working memory task (or a task thought to have a considerable working memory component). These papers include all of the miscellaneous visual working memory tasks that we could find (Visual N-back (letters), Memory-Guided Saccade, Delayed Match to Sample, Sternberg Task, Spatial Span, and the CANTAB Spatial Working Memory Task). Some prior work, e.g., Curran et al.[48], has used tasks such as the Serial Sevens Task and a Rapid Visual Information Processing Task; we consider these tasks to be less conventional measures of working memory capacity, and we did not include the Serial Sevens or the Rapid Visual Information Processing Task in our search (but see [8] for more).

**Table S3.** Literature summary of other visual working memory and working memory tasks.

| Study | Ref. | n | Route | Condition - Dose | Task | Condition - Task | p-value |
|---|---|---|---|---|---|---|---|
| Kollins et al. 2015 | [49] | 16 | Oral | 10 mg | Visual N-Back (Letters) | 0-back | n.s. |
| | | | | | | 1-back | n.s. |
| | | | | | | 2-back | n.s. |
| | | | | | | 3-back | n.s. |
| Gilman et al. 2019 | [50] | 54 | Oral | 5 – 50 mg (M = 37.8 mg) | Visual N-back (Letters) | Main effect of 0-, and 2- back | n.s., .39 |
| Ploner et al. 2002 | [51] | 12 | Oral | 10 mg | Memory Guided Saccade | N/A | * <.05 |
| D'Souza et al. 2004 | [52] | 22 | Intravenous | 2.5 mg or 5 mg | Delayed Match to Sample | 1-item, "simple shape" | 0.02 |
| | | | | | | 1-item, "complex shape" | n.s., 0.28 |
| Lane et al. 2005 | [53] | 5 | Cigarette | 2.2% THC (~18 mg) | Delayed Match to Sample | 1-item color precision | *** < .001 |
| | | | | 3.6% THC (~29 mg) | | | *** < .001 |
| Makela et al. 2006 | [54] | 19 | Oral | 5 mg | Spatial Span | Forward | n.s. |
| Foltin et al. 1993 | [55] | 7 | Cigarette | 27 mg | Sternberg Task | Set Sizes 1 – 6 | n.s. |
| Bossong et al. 2012 | [56] | 17 | Vapor | 6 mg, then 3 doses of 1 mg (30 min interval) | Sternberg Task | Set Size 1 | n.s. |
| | | | | | | Set Size 3 | * <.05 |
| | | | | | | Set Size 5 | * <.05 |
| | | | | | | Set Size 7 | n.s., <.10 |
| | | | | | | Set Size 9 | n.s. |
| Theunissen et al. 2015 | [57] | 15 | Vapor | .138 mg/kg (M = ~9.3 mg) | Sternberg Task | Set Sizes 1, 2, & 4 | n.s. |
| Ranganathan et al. 2019 | [58] | 74 | Intravenous | .05 mg/kg | CANTAB Spatial Working Memory Task | Total errors combined for set sizes 4+ (other set sizes not reported) | *** < .001 |

**Table S4.** Values for heart rate and subjective measure in Exp 1 and Exp 2. Scores are change scores for at the time of the working memory test relative to baseline. Bonferonni corrections are done within measure (3 comparisons for heart rate + blood pressure; 5 comparisons for DEQ).

| | Exp 1 | | | Exp 2 | | | |
|---|---|---|---|---|---|---|---|
| | Placebo | High (15 mg) | $p_{uncorrected}$ ($p_{bonferroni}$) | Placebo | Low (7.5 mg) | High (15 mg) | $p_{uncorrected}$ ($p_{bonferroni}$) |
| **Physiological** | | | | | | | |
| Heart Rate | **-7.77 (9.25)** | **0.73 (11.50)** | **.002 (.006)** | **-8.22 (7.27)** | **-4.57 (12.42)** | **6.48 (12.59)** | **<.001 (<.001)** |
| Systolic Blood Pressure | -2.46 (10.92) | 0.55 (7.97) | .31 (.93) | -3.04 (9.44) | -1.57 (9.97) | -2.13 (11.95) | .875 (1.0) |
| Diastolic Blood Pressure | -1.86 (8.04) | -0.09 (9.25) | .53 (1.0) | 0.04 (9.44) | 1.83 (15.82) | -2.00 (9.32) | .51 (1.0) |
| **ARCI** | **.41 (1.47)** | **4.73 (3.45)** | **<.001** | **.27 (.83)** | **2.18 (2.72)** | **3.59 (2.89)** | **<.001** |
| **DEQ** | | | | | | | |
| Feel | **4.55 (7.45)** | **45.73 (30.96)** | **<.001 (<.001)** | **2.86 (6.97)** | **24.41 (30.14)** | **37.23 (30.28)** | **<.001 (<.001)** |
| Like | **11.41 (21.73)** | **34.05 (31.37)** | **.001 (.005)** | **11.82 (24.42)** | **34.86 (28.58)** | **30.64 (27.99)** | **<.001 (<.001)** |
| Dislike | **8.73 (18.06)** | **26.09 (26.45)** | **.006 (.03)** | **6.32 (14.30)** | **16.50 (23.20)** | **33.82 (32.63)** | **.002 (.01)** |
| High | **3.05 (8.57)** | **41.91 (30.47)** | **<.001 (<.001)** | **1.82 (4.99)** | **26.09 (31.11)** | **38.82 (31.71)** | **<.001 (<.001)** |
| More | **7.55 (17.79)** | **20.64 (21.80)** | **.002 (.01)** | 10.09 (22.24) | 18.36 (24.39) | 14.95 (23.58) | .197 (.985) |

**Table S5.** Values for the Visual Analog Scales (VAS) in Exp 1. Scores are change scores for at the time of the working memory test relative to baseline. Bonferroni corrections are for the 13 comparisons in the VAS measure.

| VAS | Placebo | High (15 mg) | $p_{uncorrected}$ | $p_{bonferroni}$ |
|---|---|---|---|---|
| Anxious | -7.78 (31.26) | 3.59 (13.19) | .15 | 1.0 |
| Stimulated | -8.73 (26.85) | 15.32 (27.92) | .02 | .26 |
| Sedated | -3.91 (35.16) | -4.14 (25.42) | .98 | 1.0 |
| Elated | -12.68 (25.83) | 7.86 (28.63) | .05 | .65 |
| Insightful | -10.91 (28.29) | 11.27 (28.53) | .04 | .52 |
| **Sociable** | **-12.86 (19.12)** | **9.77 (22.51)** | **.001** | **.013** |
| Confident | -9.41 (23.00) | 6.91 (22.75) | .02 | .26 |
| Lonely | -3.09 (23.73) | 1.59 (10.45) | .44 | 1.0 |
| Playful | -11.82 (21.41) | 14.96 (26.51) | .004 | .052 |
| Dizzy | -6.86 (25.17) | 4.73 (23.66) | .17 | 1.0 |
| Loving | -5.09 (31.03) | 6.27 (24.83) | .24 | 1.0 |
| **Friendly** | **-10.77 (27.62)** | **14.82 (24.94)** | **.003** | **.039** |
| Restless | -7.55 (38.98) | 8.27 (29.31) | .11 | 1.0 |

**Table S6.** End of Session Questionnaire (ESQ) perceived drug reports in Experiment 1 and Experiment 2. At the end of each session, participants were asked to choose from a list of options which drug they think they may have received. Each cell shows subject counts for perceived drug condition.

| | Placebo | Cannabinoid | Sedative | Stimulant | Not Sure |
|---|---|---|---|---|---|
| **Exp. 1:** Placebo Session | 17 | 0 | 6 | 1 | 0 |
| **Exp. 1:** Drug Session (15 mg) | 0 | 17 | 4 | 3 | 0 |
| **Exp. 2:** Placebo Session | 15 | 2 | 5 | 2 | 0 |
| **Exp. 2:** Drug Session (7.5 mg) | 4 | 12 | 4 | 4 | 0 |
| **Exp. 2:** Drug Session (15 mg) | 2 | 16 | 5 | 0 | 1 |
